# Motor Imagery EEG neurofeedback skill acquisition in the context of declarative interference and sleep

**DOI:** 10.1101/2020.12.11.420919

**Authors:** Mareike Daeglau, Catharina Zich, Julius Welzel, Samira Kristina Saak, Jannik Florian Scheffels, Cornelia Kranczioch

## Abstract

Motor imagery (MI) practice in combination with neurofeedback (NF) is a promising supplement to facilitate the acquisition of motor abilities and the recovery of impaired motor abilities following brain injuries. However, the ability to control MI NF is subject to a wide range of inter-individual variability. A substantial number of users experience difficulties in achieving good results, which compromises their chances to benefit from MI NF in a learning or rehabilitation context. It has been suggested that context factors, that is, factors outside the actual motor task, can explain individual differences in motor skill acquisition. Retrospective declarative interference and sleep have already been identified as critical factors for motor execution (ME) and MI based practice. Here, we investigate whether these findings generalize to MI NF practice.

Three groups underwent three blocks of MI NF practice each on two subsequent days. In two of the groups, MI NF blocks were followed by either immediate or delayed declarative memory tasks. The control group performed only MI NF and no specific interference tasks. Two of the MI NF blocks were run on the first day of the experiment, the third in the morning of the second day. Significant within-block NF gains in mu and beta frequency event-related desynchronization (ERD) where evident for all groups. However, effects of sleep on MI NF ERD were not found. Data did also not indicate an impact of immediate or delayed declarative interference on MI NF ERD.

Our results indicate that effects of sleep and declarative interference context on ME or MI practice cannot unconditionally be generalized to MI NF skill acquisition. The findings are discussed in the context of variable experimental task designs, inter-individual differences, and performance measures.

## 1. Introduction

The acquisition of new movements or improving existing motor skills is a significant part of everyday life. It is known that motor acquisition is mainly achieved by repeatedly physically executing the target movements, incorporating the external sensory input, and adapting subsequent movement attempts (Adams, 1971; Hikosaka, Nakamura, Sakai, & Nakahara, 2002; Willingham, 1998). In addition to this motor execution (ME) practice loop, motor acquisition can be supported by motor imagery (MI) (e.g. Guillot, Moschberger, & Collet, 2013; Ruffino, Papaxanthis, & Lebon, 2017). MI is a dynamic mental state, which involves a systematic internal simulation process to rehearse a target movement without overtly executing it (Decety, 1996; Di Rienzo et al., 2016). The neural simulation of action theory provides a theoretical framework for the interplay between ME and MI (Jeannerod, 2001). It postulates that imagined movements are functionally equivalent to executed ones not only in terms of overt motor stages, as e.g. motor planning, but also with regard to the underlying neural networks (for a critical review, see O’Shea & Moran, 2017). Remarkably, the activation of these specific networks strongly depends on the MI strategy, i.e., the activity of sensorimotor areas is predominantly induced by kinaesthetic MI (internal perspective) and less by visual MI (external perspective) (Annett, 1995; Hétu et al., 2013; Neuper, Scherer, Reiner, & Pfurtscheller, 2005; Stinear, Byblow, Steyvers, Levin, & Swinnen, 2006).

MI practice is suitable to facilitate the acquisition of new motor skills and the improvement of already existing motor skills in healthy subjects (Kraeutner, McWhinney, Solomon, Dithurbide, & Boe, 2018; Schack, Essig, Frank, & Koester, 2014; Wriessnegger, Brunner, & Müller-Putz, 2018). Furthermore, MI practice is considered as a promising tool to complement motor rehabilitation interventions after a brain injury, such as stroke, by facilitating the relearning of lost motor skills (Bajaj, Butler, Drake, & Dhamala, 2015; Maier, Ballester, & Verschure, 2019; Malouin, Jackson, & Richards, 2013). To overcome the lack of sensory feedback in MI and so to enhance its positive effects, MI can be combined with neurofeedback (NF).

NF serves as a channel for feeding back information about MI performance to the subject, and can be utilized to enhance adaptive activation patterns (Braun, Emkes, Thorne, & Debener, 2016; Ietswaart et al., 2011; Zich, De Vos, Kranczioch, & Debener, 2015), motor skill acquisition, and motor recovery (for a review, see Pichiorri & Mattia, 2020). MI NF paradigms are typically controlled by event-related (de)synchronization (ERD/S), reflecting task related power changes of rhythmic brain activity recorded within the mu (8-12 Hz) and beta (13-30 Hz) frequency range over sensorimotor areas contralateral to the target limb (Cheyne, 2013; Lopes da Silva & Pfurtscheller, 1999; Pfurtscheller & Aranibar, 1979).

The active self-regulation of MI induced ERD/S and the closely interrelated MI NF control can be considered as skills that can be acquired and that are subject to principles of learning (Lotte et al., 2015; Lotte, Larrue, & Mühl, 2013). For ME and MI skill acquisition it has been shown that they are influenced by task context (Brown & Robertson, 2007; Debarnot et al., 2012; Debarnot, Castellani, Valenza, Sebastiani, & Guillot, 2011; Schlatter et al., 2020). Based on this it can be expected that for achieving MI NF control, the context in which practice takes place matters (e.g. Daeglau et al., 2020; Roc, Pillette, N’Kaoua, & Lotte, 2019).

Two relevant context factors for MI skill acquisition are declarative interference through a subsequent non-motor task and sleep (e.g. Brown & Robertson, 2007; Debarnot et al., 2012; Debarnot, Castellani, et al., 2011). Debarnot and colleagues (2012) demonstrated for a finger tapping task that physical task performance was negatively affected by declarative interference following MI practice both over intervals of sleep and wakefulness. Notably, this contrasts with ME practice, where declarative interference similarly impairs the consolidation of a motor task over wakefulness, whereas sleep has been shown to support performance recovery (Brown & Robertson, 2007; but see, Rothkirch, Wolff, Margraf, Pedersen, & Witt, 2018).

Studies on the role of sleep in ME motor skill acquisition and without specific interference tasks have indicated that sleep following ME skill acquisition leads to additional gains in the practised motor skill (for a review, see King, Hoedlmoser, Hirschauer, Dolfen, & Albouy, 2017). Other studies however challenge this notion (Brawn, Fenn, Nusbaum, & Margoliash, 2010; Hotermans, Peigneux, De Noordhout, Moonen, & Maquet, 2006; Nettersheim, Hallschmid, Born, & Diekelmann, 2015) Using a finger-tapping task, Nettersheim and colleagues (2015) showed that the supposedly sleep related gain is not really a gain but rather a stabilisation of task performance at the early boost level. The early boost describes an offline gain in performance that can be measured around 5-30 minutes after the motor task and then decays over the next 4-12 hours of wakefulness (Brawn et al., 2010; Hotermans et al., 2006). Gains in performance following a sleep period have also been reported for MI practice (Debarnot, Clerget, & Olivier, 2011; Debarnot, Creveaux, Collet, Doyon, & Guillot, 2009; Debarnot, Creveaux, Collet, Gemignani, et al., 2009). This has been interpreted as follows: motor consolidation and associated delayed offline gains in ME performance acquired through MI practice profit from sleep or even depend on it. Yet it seems possible that this interpretation needs to be reconsidered, and that also for MI skill acquisition sleep does not result in an additional gain but rather in a consolidation at the early boost level (Nettersheim et al., 2015). Supporting this assumption, an early boost can also be found for MI skill acquisition (Debarnot, Clerget, et al., 2011).

To date, it has not been studied how sleep and declarative interference affect MI NF skill acquisition, and whether MI NF skill acquisition is subject to an early boost and subsequent decay effects. To address these questions, three groups underwent three MI NF blocks each, which took place over two consecutive days. In the two experimental groups, two of the tree MI NF blocks were arranged thus that each MI NF block was followed by declarative interference tasks (immediate interference) or thus that the declarative interference tasks followed after a day of wakefulness (late interference). In the control group MI NF blocks were combined with control tasks and no explicit declarative interference task was performed (no interference). MI (NF) ERD of the MI NF blocks was used as a quantifiable feature for describing MI (NF) performance related to MI (NF) skill acquisition. Regarding immediate declarative interference, we hypothesised it to have an adverse impact on MI NF performance compared to no interference, as evident in reduced contralateral ERD within the mu and beta frequency range in the group receiving immediate declarative interference. We additionally hypothesized that sleep would not protect MI NF performance from these adverse effects of declarative interference. Further, regarding late interference, we expected to observe an early boost in MI NF performance from the first to the second MI NF block that would not be affected by late interference but rather consolidated after a night of sleep.

## 2. Material and Methods

### 2.1 Participants

We collected data from 66 healthy, young participants. All participants reported normal or corrected-to-normal vision. None of the participants reported a current or previous history of psychiatric or neurological conditions or use of psychoactive medication. As indicated by the Edinburgh Handedness Inventory (Oldfield, 1971), all participants were right-handed. Participants did not participate in previous MI NF studies. Explicit information about the purpose of the conducted experiments were not provided. Participants were informed about MI processes in general and specifically about the characteristics of visual and kinaesthetic MI. Every participant read and signed an institutionally approved consent form prior to the experiment. A total of 13 data sets were discharged from analyses. Three data sets were discarded due to technical issues during data collection, six data sets due to participant’s early drop out, one data set due to non-compliance to task instructions (i.e. moving during MI blocks), and three data sets because post-hoc analysis of self-report questionnaires indicated that participants did not comply with the general instruction to stay drug free between the three experimental session.

The final sample consisted of 53 participants with three sessions each. Final group sizes were 17 participants in *group late-interference* (14 women, aged 20 – 32 years, *M* and *SD*: 24.3 ± 3.5 years), 19 participants in *group immediate-interference* (17 women, aged 21 – 35 years, *M* and *SD*: 25.1 ± 3.9 years) and 17 participants in *group no-interference* (10 women, aged 23 – 32 years, *M* and *SD*: 25.8 ± 2.5 years).

The study protocol was approved by the Commission for Research Impact Assessment and Ethics of the University of Oldenburg.

### 2.2 Study Layout

The study design is illustrated in Figure 1A. The study consisted of three experimental sessions spread across two successive days. Two sessions were run in the morning of the first and the second day (8:30 – 12:00), and one in the evening (18:00 – 21:30) of the first day. Sessions consisted of a combination of three out of four different blocks (motor imagery, interference, control, break). Motor imagery or MI blocks consisted of one run MI without NF and two MI NF runs (for details see section 3.3.1 Motor Imagery Neurofeedback Paradigm). Interference blocks encompassed four different cognitive demanding non-motor tasks (for details see section 3.3.2 Interference-, Break- and Control-Blocks). In control blocks and break blocks participants passively watched a nature documentary. Each block lasted about 30 minutes.

**Fig. 1:**
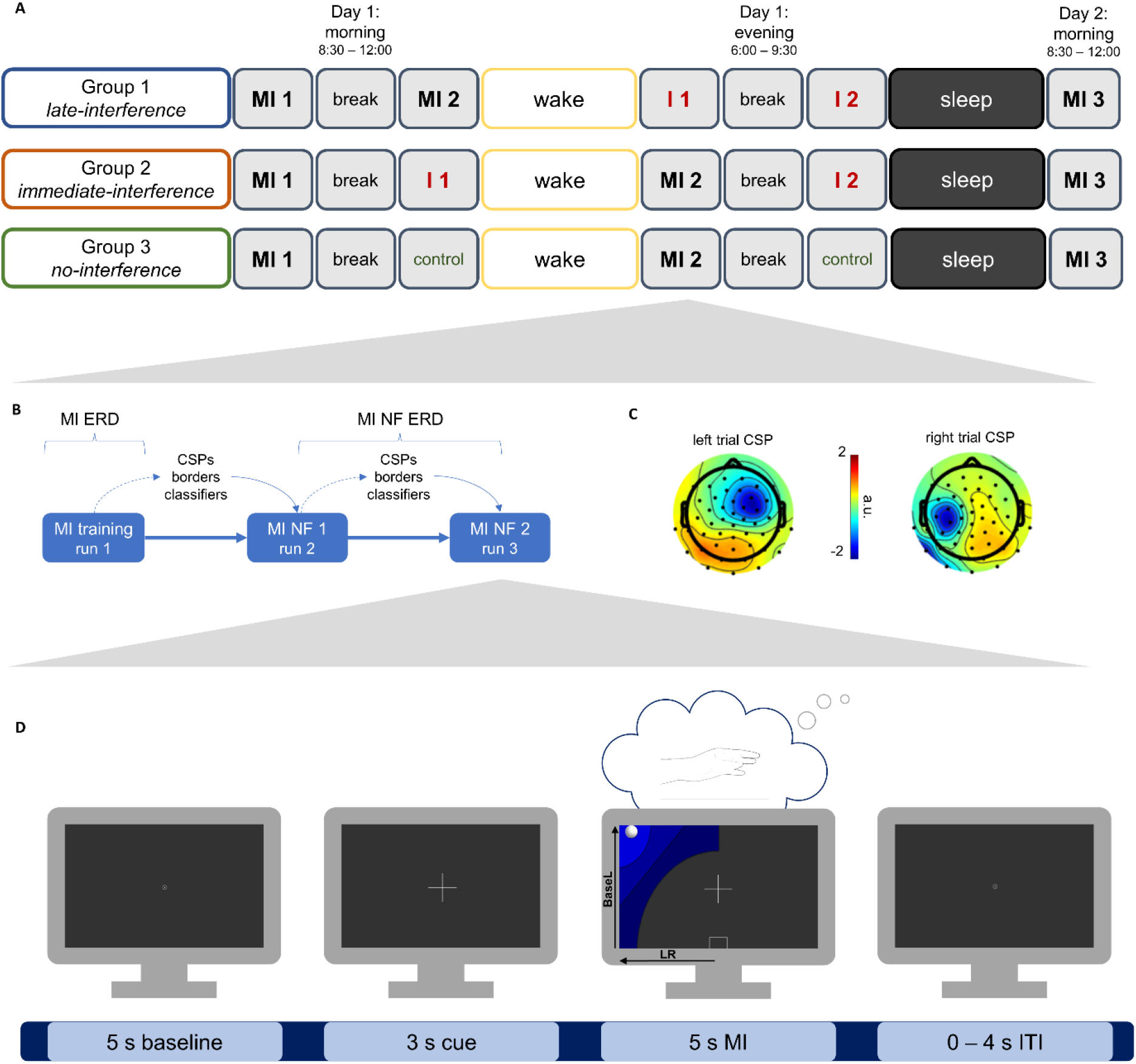
Experimental Hierarchy. **A. Study Design**. The first session in the morning of day 1 began with a block of MI NF practice (**MI1**) followed by a **break** of 30 minutes (watching a documentary). Afterwards *group late-interference* performed another block of MI NF practice (**MI2**), *group immediate-interference* completed a block of declarative non-motor interference (**I1**) and *group no-interference* continued watching the documentary as a **control** condition. Participants then followed their daily life routine. However, daytime naps or excessive sport activities were not allowed, which was monitored by an activity tracker. In the evening of the same day, *group late-interference* completed its first interference block (**I1**), while *group immediate-interference* and *group no-interference* completed their second MI NF block (**MI2**). After a **break** *group late-interference* and *group immediate-interference* had their second interference block (**I2**) and *group no-interference* proceeded with its **control** condition. After a night of sleep all groups returned for a final session with a single block of MI NF practice (**MI3**). **B. Flowchart MI NF block.** Each MI block encompassed three runs consisting of 40 trials each (20 left hand, 20 hand right trials). After the training run, where no feedback was provided, NF parameters common spatial patterns (CSPs), classifier, and border were calculated and set for the second run (MI NF1). After this first feedback run (MI NF1), the NF parameters were renewed and set for the third run (MI NF2). For offline evaluation, the event-related-desynchronization (ERD) was calculated from EEG data obtained from MI NF 1 and MI NF 2 and averaged (MI NF ERD). **C. Representative single subject CSPs**. CSPs shown are based on left hand and right-hand trials of one MI run. **D. Structure of a left-hand motor imagery (MI) neurofeedback (NF) trial.** Each trial began with a baseline period of 5 s showing the outline of a small circle. Afterwards a fixation cross displayed for 3 s indicated the imminent start of the MI task period. The onset of a graphic comprising 3 different shades of blue indicated the beginning of the task period (duration 5 s). The location of the graphic indicated which hand to use for MI. During the NF runs a white ball moved along the horizontal (LR for left vs. right MI) and vertical (BaseL for baseline vs. left MI) axes according to the classifier output magnitudes. In the training runs the ball remained motionless in the centre of the screen. The inter-trial-interval (ITI) comprised 0-4 s connecting to the next baseline period

Participants were assigned in alternation on a rolling basis to either *group late-interference*, *group immediate-interference* or *group no-interference*. The first experimental session of *group late-interference* comprised two blocks of MI NF (MI1, MI2), separated by a break block. For *group immediate-interference* it consisted of a block of MI NF (MI1), a break block, and an interference block (I1), and for *group no-interference* of one block of MI NF (MI1), a break block and a control block. Participants were instructed to follow their regular daily routine in-between sessions, except for taking naps and exhausting sport activities. Adherence to these instructions was monitored using an actigraphy watch (MotionWatch8, CamNtech Ltd., Cambridgeshire, UK). The second experimental session was performed in the evening of the first day. For *group late-interference* it covered two blocks of interference (I1, I2) and a break block in-between, for *group immediate-interference* one block of MI NF (MI2), a break block and an interference block (I2) and for *group no-interference* one block of MI NF (MI2), a break block and a control block. Participants were instructed to get at least 8 hours of sleep, which was confirmed by the actigraphy watch. The third experimental session was conducted the next morning and consisted of a single block of MI NF (MI3) for all three groups.

In both morning sessions, participants completed the Stanford Sleepiness Score (SSS; Hoddes, Zarcone, & Dement, 1972) questionnaire, which provides a subjective measure of alertness. The SSS is a 7- point scale, with 1 being the most alert state. The SSS was applied to ensure an adequate state of alertness prior to each session for all participants. For each of the three sessions, participants had to rate their current alertness at least a 4 (“Somewhat foggy, let down”) but not below for their dataset to be included in analysis. Prior to the first experimental session the short version of the kinaesthetic and visual imagery questionnaire (KVIQ; Malouin et al., 2007) was conducted to emphasize the difference between visual and kinaesthetic MI. After MI NF blocks participants rated their motivation on a visual analogue scale, and perceived vividness and easiness of MI on a 5-point Likert scale. Participants further completed the Pittsburg Sleep Quality Index to assess sleep quality and quantity (PSQI; Buysse, Reynolds, Monk, Berman, & Kupfer, 1989) and the Epworth Sleepiness Scale (ESS; Johns, 1991) to assess their ‘daytime sleepiness’. The PSQI examines retrospectively over a period of four weeks about the frequency of sleep disturbing events, the assessment of sleep quality, the usual sleeping times, sleep latency and duration, the intake of sleep medication and daytime sleepiness. A total of 18 items are used for quantitative evaluation and assigned to seven components, each of which ranges from 0 to 3. The total score is the sum of the component scores and can vary from 0 to 21, whereby a higher score corresponds to a lower quality of sleep. The ESS is a self-administered questionnaire covering eight questions. Participants had to rate their usual chances of dozing off or falling asleep while performing eight different activities (4-point scale: 0-3). The ESS score ranges from 0 to 24. The higher the ESS score, the higher that person’s average sleep propensity in daily life, or their ‘daytime sleepiness’. As part of a different research question, participants also performed a limb lateralization task (LLT; Ter Horst, Van Lier, & Steenbergen, 2010) and the nine-hole-peg test (NHPT; Kellor, Frost, Silberberg, Iversen, & Cummings, 1971).

Participants were instructed to be free of drug, alcohol, and caffeine for 24 h prior to and during each experimental session.

### 2.3 Experimental Procedure

#### 2.3.1 Motor Imagery Neurofeedback Paradigm

A previously established MI NF paradigm (Braun et al., 2017; Meekes, Debener, Zich, Bleichner, & Kranczioch, 2019; Zich, Debener, Schweinitz, et al., 2017) was adapted for the present study. In the present implementation, in each MI block participants performed one run of MI without NF followed by two runs of MI NF (cf. Fig. 1B). Each run had a duration of about ten minutes. The imagined movement was sequential thumb to finger opposition either with the left or the right hand starting with the little finger. Prior to the first run, the movement was demonstrated by the experimenter. The participant was verbally instructed to copy the movement and practice it with both hands until they felt sufficiently familiarized. Movements were performed at a rate of about 1 Hz. Participants were instructed to hold this pace during kinaesthetic MI. In the first MI run of each block no NF was presented. The EEG data recorded in this calibration/training run were used to set up the NF parameters for the NF in the second MI run. In turn, EEG data acquired in the second MI run served for calibrating the parameters used for the NF in the third MI run. Each MI run consisted of 20 left and 20 right hand trials presented in pseudo-randomized order.

Stimulus presentation was controlled with OpenViBE 0.17.1 (Renard, Congedo, Delannoy, & Le, 2010). The NF was based on the adapted Graz MI protocol as implemented in OpenViBE (Renard et al., 2010; Zich, Debener, De Vos, et al., 2015). Each trial began with a baseline period of 5 s showing a small, outlined circle. The circle was replaced by a fixation cross displayed for 3 s, indicating the imminent start of the MI task period. The start of the MI task period was signalled by a blue graphic displayed in addition to the fixation cross. The blue graphic was placed either on the left or right half of the screen (see Fig. 1D. The on-screen location of the blue graphic signalled the hand to be used during the MI task period. Each task period had a duration of 5 s. In the last two MI blocks NF was included in the task period. The NF was visualized as a white ball moving along two dimensions on either the left or right half of the screen. The horizontal position of the ball reflected the degree of ERD lateralization, the vertical position the degree of contralateral ERD (see Fig.1D). The exact horizontal and vertical positions of the ball were determined by the output of two classifiers reflecting the difference between contralateral and ipsilateral EEG activity and between baseline and contralateral EEG activity (see section EEG analysis for details). Participants were informed that navigating and maintaining the ball in the upper left or right corner of the screen, depending on the location of the graphic, reflected an appropriate task performance. The NF screen was updated at a frequency of 16 Hz. During the inter-trial interval, the screen showed a small, outlined circle presented pseudo-randomly for 0 to 4 s (increments of 1 s). Participants were instructed to sit still but relaxed. NF borders, representing the maximum reachable edges on screen for both the vertical and horizontal dimensions of the NF, were kept constant within a run and defined as the upper quartile of the classifier output from the previous run.

#### 2.3.2 Interference-, Break- and Control-Blocks

Declarative interference blocks comprised four different non-motor tasks: a word list recall, an n-back task, a face-name matching task, and a modified version of the Paced Auditory Serial Addition Test (PASAT; Gronwall, 1977).

For the word list recall task, lists of 24 words were presented visually three times. Words were presented for 3 s without an ITI. Participants were instructed to remember as many words as possible. After a break of 15 – 20 minutes, participants had to identify remembered words among distractor words via a button press as fast and accurate as possible. Wordlists were retrieved from Salvidegoitia and colleagues (Piñeyro Salvidegoitia et al., 2019; CELEX online lexical database, Max Planck Institute for Psycholinguistics, 2001, available at http://celex.mpi.nl/).

The n-back task consisted of one training block (eight trials) to familiarize participants with the task followed by two blocks (120 trials). Participants had to indicate whether the current letter was identical to the previous one (1-back task) as fast and accurate as possible. Each letter was presented for 600 ms.

The face-name matching task was designed based on the corresponding subtest from the memory assessment scales (MAS; Williams, Williams, & Gillard, 1991). The procedure started with a learning phase, where participants were successively presented 20 faces in combination with a name (each 5 sec). They were asked to remember each name-face combination. Thereafter, they were shown one of the 20 faces at once and three names at the bottom, one being the correct name and two previously learned but incorrect names. Participants were asked to indicate the location of the correct name via a button press on the keyboard. This procedure was repeated for all 20 faces. Images for the faces were retrieved from pics.stir.ac.uk.

The PASAT was conducted following the instructions in the manual (Gronwall, 1977) except that digits were not presented verbally but visually for 2 s each without ISI. A total of 122 digits were presented in two blocks. Participants were asked to add up the last two digits and say the result out loud. As soon as the next number was shown, it had to be added to the previous one again.

Each interference task was performed once per interference block. To minimize task familiarization effects across sessions various versions were created for all interference tasks. Interference tasks were presented in pseudo-randomized order across participants within each interference block. Results were recorded but are not reported here.

Break and control blocks consisted of 30-minute sections of various nature documentaries each with male narrators and ambient music, but no visible human interaction. The same set of documentaries was shown to all participants, but each documentary was shown to each subject only once. All tasks were controlled by customized scripts implemented in Presentation software (Version 17.0, Neurobehavioral Systems, Inc., Berkeley, CA, USA, RRID:SCR_002521).

### 2.4 Data Acquisition

EEG data were acquired from 65 sintered Ag/AgCl electrodes using an equidistant infracerebral electrode layout with a central frontopolar site as ground and a nose tip reference (Easycap, Herrsching, Germany). In addition, bipolar surface EMG was recorded from both hands and arms by placing sintered Ag/AgCl electrodes over the muscle belly and the proximal base of the Flexor digitorum superficialis and the Abductor pollicis longus with reference and ground on the collarbone. Both EEG and EMG data were recorded using a BrainAmp amplifier system (Brain-Products, Gilching, Germany). Data were obtained with an amplitude resolution of 0.1 μV and a sampling rate of 500 Hz with online analogue filter settings of 0.016 to 250 Hz. Electrode impedances were maintained below 10 kΩ for the EEG and below 100 kΩ for the EMG before data acquisition. Data acquisition was performed using the OpenViBE acquisition server 0.17.1 [29]. In addition, resting state EEG recordings of two minutes each were obtained before and after each session using BrainVision recorder software (Version 1.20.0506, Brain-Products GmbH, Gilching, Germany).

### 2.5 Data Analysis

#### 2.5.1 Online Processing

Online EEG data analyses for providing NF comprised three parts and was performed after the first and second MI run (i.e. training/calibration and MI NF1). In the first and second part of the analyses subject-specific parameters for the subsequent MI NF blocks were determined by means of common spatial pattern (CSP) analysis in MATLAB (Version 9.3; MathWorks, Natick, MA, USA, RRID:SCR_001622), and classifier training and border computation in OpenViBE (Renard et al., 2010). The third part encompassed the actual NF delivery during the second and third experimental runs through OpenViBE using the results of the previous parameter estimation.

For the CSP analysis, EEG data from the central 49 channels were high-pass filtered at 8 Hz (finite impulse response, filter order 826) and subsequently low-pass filtered at 30 Hz (finite impulse response, filter order 220) using EEGLAB toolbox Version 14.1.1 (Delorme and Makeig, 2004) for MATLAB (Version 9.3; MathWorks, Natick, MA, USA, RRID:SCR_001622). This filter range was set to encompass the sensorimotor rhythms mu (8–12 Hz) and beta (13–30 Hz), to which the neural correlate of interest, the event-related desynchronization or ERD, is highly specific to [18,19]. Epochs were extracted from 0.5 to 4.5 s relative to MI onset, separately for left- and right-hand trials. Segments containing artifacts were rejected (EEGLAB function pop_jointprob.m, SD = 3) and the remaining data submitted to a CSP analysis pipeline (Ramoser et al., 2000). CSP analysis is a common approach to obtain spatial filters optimized for the detection of power differences between two classes (i.e. left vs right hand MI) by maximizing the variance of the signal for one class (i.e. left hand MI) while simultaneously minimizing the variance of the signal for a second class (i.e. right hand MI). For each class, of the three filters with the highest variance segregation for the class the most neurophysiologically plausible filter was selected, and the filter coefficients of the two selected CSPs (one for each class) were submitted to OpenVibe. Exemplary single subject CSPs for one MI NF run are shown in Fig. 1C.

For the classifier training in OpenVibe, EEG raw data were spatially filtered using the selected CSPs and temporally filtered using a 4th-order Butterworth band-pass filter (8 – 30Hz, 0.5 dB pass band ripple). Epoching in left- and right-hand MI periods was done from 0.5– 4.5 s relative to the onset of MI and, also relative to MI onset from −7 to −3 s for the corresponding baseline intervals. The resulting intervals were subdivided into 1 s time bins overlapping by 0.9375 s. The logarithmic power of the band-pass-filtered 1 s time windows represented the features for linear discriminant analysis (LDA) classification using sevenfold cross-validation (Fisher, 1936). For the online NF, two classifiers per active side were trained: either left motor imagery vs. baseline (BaseL), or right motor imagery vs. baseline (BaseR), representing the vertical component of the feedback (contralateral ERD) and left motor imagery vs. right motor imagery (LR) controlling the horizontal component of the feedback (degree of lateralization). Based on the results of the cross-validation three border values were calculated, corresponding to the upper quartiles of the three classification distributions. These border values were used to determine the range of the display for the horizontal and vertical axis for the online NF. CSPs, classifiers and borders were updated for the second MI NF run of a MI NF block based on data of the first MI NF run. Obtained parameters were applied to the respective successive run.

Online classification accuracies (*M* ± *SD*) were for *group late-interference* MI NF block 1 69.2 ± 6.3%, MI NF block 2 69.4 ± 5.7% and MI NF block 3 70.6 ± 6.2%, for *group immediate-interference* MI NF block 1 70.3 ± 8.0%, MI NF block 2 70.2 ± 6.4% and MI NF block 3 70.1 ± 7.3 %, and for *group no-interference* MI NF block 1 66.3 ± 6.4%, MI NF block 2 69.6 ± 5.9 and MI NF block 3 67.4 ± 6.1%. With 80 trials in total (40 per MI NF run), online classification accuracies were significantly above the chance level of 58.7% (Combrisson & Jerbi, 2015).

#### 2.5.2 Offline Analysis

##### 2.5.2.1 EMG Analysis

EMG data were filtered with a cut-off frequency of 25 Hz using a high-pass finite-impulse response filter with a hamming window (filter order: 264). Noise removal was conducted via wavelet denoising (wavelet signal denoiser toolbox, MathWorks, Natick, MA, USA) with a Daubechies 4 (dB4) wavelet. EMG data were then segmented from −9 s relative to +7 s. For each trial, the standard deviation and the 250-samples centred moving standard deviation were calculated. Trials in which the moving standard deviation exceeded the standard deviation of the trial by the factor 2.5 at any point were considered to contain movement artifacts and excluded from further analyses (M and SD: 49 ± 35.6 trials, range: 1-141 trials for a total of 360 trials).

##### 2.5.2.2 EEG Analysis

EEG data were preprocessed with the EEGLAB toolbox Version 14.1.1 (Delorme & Makeig, 2004) for MATLAB (Version 9.3; MathWorks, Natick, MA, USA). For artifact correction independent components analysis (ICA) was performed. The EEG data of all three runs within one experimental block, i.e. training, MI NF1 and MI NF2, were appended for further processing. Identification of improbable channels was conducted using the EEGLAB extension trimOutlier (https://sccn.ucsd.edu/wiki/EEGLAB_Extensions) with an upper and lower boundary of two standard deviations of the mean standard deviation across all channels (channels identified M and SD: 1.9 ± 0 channels, range 0 – 4 channels). Channels exceeding this threshold were removed accordingly. A copy of the data was first low-pass filtered (40 Hz, FIR, hamming window, filter order: 166), down-sampled to 250 Hz and high-pass filtered (1 Hz, FIR, hamming window, filter order: 414). Afterwards data were segmented into consecutive one-second epochs and segments containing artefacts were removed (EEGLAB functions pop_jointprob.m, pop_rejkurt.m, both SD = 3). Remaining data were submitted to the extended infomax ICA to estimate the unmixing weights of 45 independent components. The unmixing matrix obtained from this procedure was applied to the original unfiltered EEG dataset for selection and rejection of components representing stereotypical artifacts. Components reflecting eye, muscle and heart activity were identified using ICLabel (Pion-Tonachini, Kreutz-Delgado, & Makeig, 2019), and by the Eye-Catch approach (Bigdely-Shamlo, Kreutz-Delgado, Kothe, & Makeig, 2013) and controlled by visual inspection. Components flagged as artifacts were removed from further analysis. Artifact corrected EEG data were low-pass filtered with a finite-impulse response filter and a cut-off frequency of 30 Hz (hamming window, filter order 220, Fs = 500 Hz), and subsequently high-pass filtered with a finite-impulse response filter and a cut-off frequency of 8 Hz (hamming window, filter order 826, Fs = 500 Hz). After the data were re-referenced to common average, and bad channel signals were replaced by spherical interpolation. Data were segmented from −7 s to 9 s relative to the start of the task interval, separately for left and right trials, and baseline corrected (−6 to −4 s). Artefactual epochs as indicated by the joint probability within each of the experimental runs (EEGLAB function pop_jointprob.m, pop_rejkurt.m, both SD = 3) and epochs flagged by the EMG analysis (see 2.5.2.1 EMG Analysis) were discarded from further analyses. In parallel to the online analysis, EEG data were reduced to the central 49 channels. CSP filters were calculated offline following the same procedure as described for the online EEG analysis, except that data based on all three MI runs was used. Likewise, two CSP filters were selected, one for left and one for right hand MI. Contralateral activity was obtained by multiplying CSP filters with corresponding trials (i.e. right CSP filters with right trials EEG data, left CSP filters with left trials EEG data) and ipsilateral activity by multiplying CSP filters with respective opposite trials (i.e. right CSP filters with left trials EEG data, left CSP filters with right trials EEG data). Thereafter, task-related event-related desynchronization (ERD) was extracted following the procedure proposed by Lopes da Silva & Pfurtscheller (1999). Since hemispheric differences were not of interest in the present study, relative ERD contralateral to the target hand was averaged for left- and right-hand trials. On average, a total of 31.1 ± 4.5 trials per subject and condition (range 24 – 40 trials) were used for analysis. For the statistical analyses, contralateral relative ERD was averaged within MI runs across a time window of 0.5 to 4.5 s with respect to the onset of the MI task interval. For the evaluation of MI NF runs, contralateral relative ERD was additionally averaged across MI NF1 and MI NF2 resulting in MI NF ERD. MI ERD (training/calibration) and MI NF ERD (NF1/NF2) serve as quantifiable features to describe MI (NF) performance related to MI (NF) skill acquisition in this study.

### 2.6 Statistical Analyses

To ensure no initial differences across groups, questionnaire data (motivation, tiredness, MI vividness/easiness, sleep scores) and MI NF ERD were analysed by means of separate Bayes ANOVAs with between-subject factor group and aforementioned data as respective dependent variable were conducted.

To confirm the effect of NF on MI ERD, we performed paired t-tests within each group and block comparing the training run MI ERD with MI ERD obtained from both MI NF runs. This was followed by a mixed 3×3-ANOVA with group as between-subject factor and block as within-subject factor to explore how the NF-related gain in MI ERD evolved over time. The dependent variable in this analysis was the difference between training run (MI ERD) and both NF runs (MI NF ERD) within one block.

Hypotheses were tested in planned comparisons. Frequentist statistics were used whenever a difference was expected, Bayes statistics were used when no difference was expected. For interference it was hypothesized that immediate declarative interference has an adverse impact on MI NF performance compared to no interference. To test whether immediate interference reduced MI NF ERD following a period of wakefulness, an independent samples t-test between MI NF block 2 of *groups immediate-interference* and *no-interference* was performed.

We further hypothesized that sleep would not eliminate the adverse effects of declarative interference in MI NF performance, meaning that performance would not be better after a night of sleep. This was tested by a Bayes paired t-test between MI NF blocks 2 and 3 of *group immediate-interference*.

For group *late-interference* we expected to observe an early boost effect in MI NF performance as well as a consolidation of this performance level after a night of sleep. This was tested in a repeated measures ANOVA comprising MI NF blocks 1 – 3 for *group late-interference*. The ANOVA was complemented by a Bayes paired t-test between MI NF block 2 and MI NF block 3.

In case that sphericity was violated Greenhouse–Geisser–correction was applied as implemented in the R-package ez (Version 4.4-0; Lawrence, 2016). Post hoc comparisons were conducted using two-tailed t-tests. Multiple pairwise comparisons were corrected for by the Holm-Bonferroni method according to the number of performed tests (Holm, 1979). All numerical values are reported as mean ± SE. Effect sizes are reported as Eta-squared (η²) for ANOVAs and Cohen’s d (d) for t-tests. If Frequentist statistics showed non-significant results, they were followed up by Bayes statistics to test the confidence in the Null hypothesis.

All Frequentist statistics were conducted as implemented in RStudio (Version 1.1.463; Team, 2018). All Bayes statistics were performed with the free software JASP using default priors (Version 0.9.2.0; JASP Team, 2019).

## 3. Results

The baseline and task period time-courses of MI NF ERD for all groups and MI NF blocks are shown in Figure 2. Clear ERD responses could be confirmed for all groups and blocks. Figure 3 shows individual MI NF ERDs split by group and MI blocks. Descriptively, at the group level, MI NF ERD did not differ significantly over MI NF blocks nor between groups. At the single subject level differences across MI NF blocks were present but showed a high variability overall and no clear within-group pattern.

**Fig. 2:**
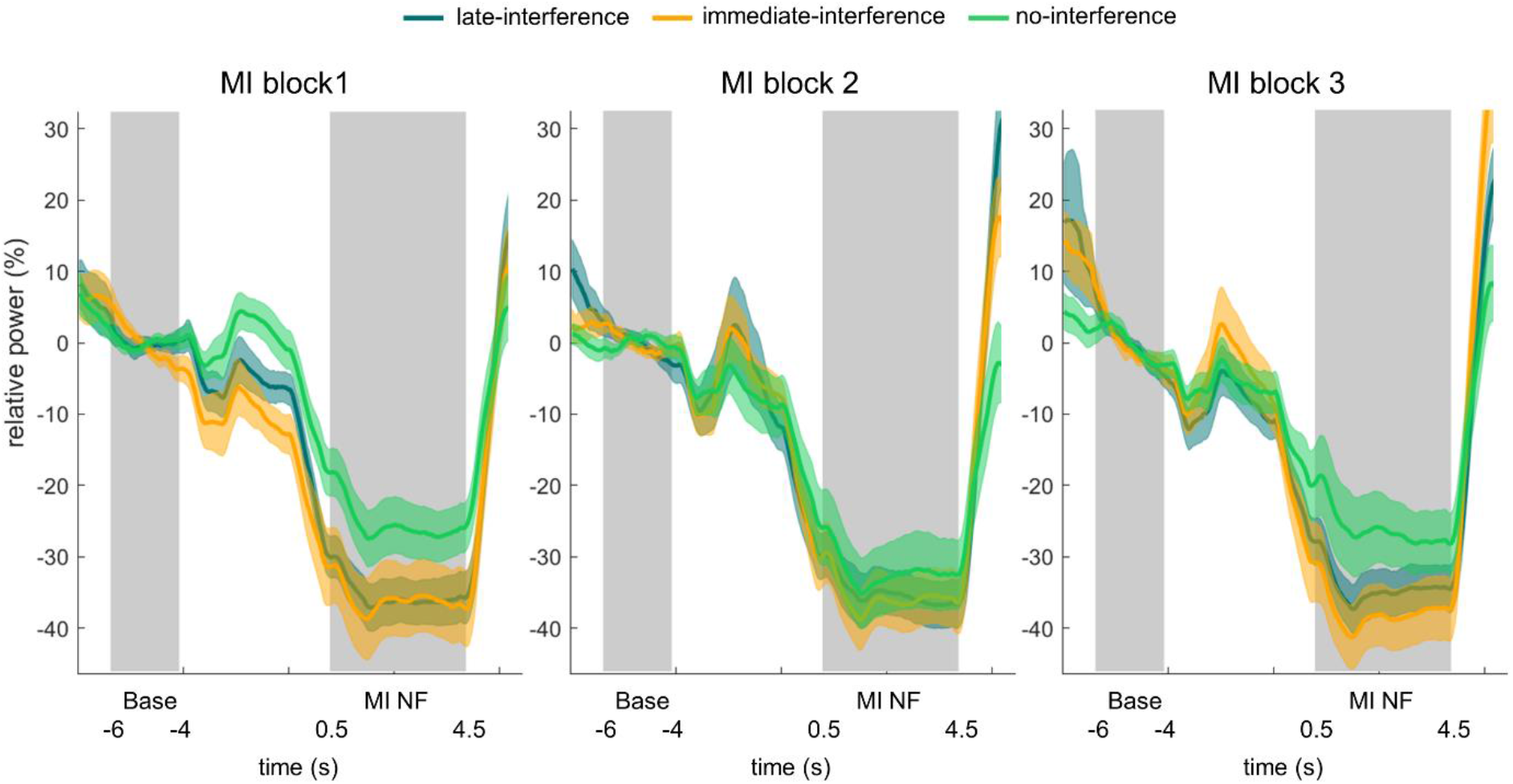
MI NF ERD time courses. MI NF ERD time courses are shown for groups *late-interference* (blue), *immediate-interference* (orange) and *no-interference* (green) for each MI block. Grey areas indicate the baseline (−6 to −4 s before task onset) and the statistically analysed MI interval (0.5 – 4.5 s after task onset). Data are averaged across MI NF runs within a block.

**Fig 3.**
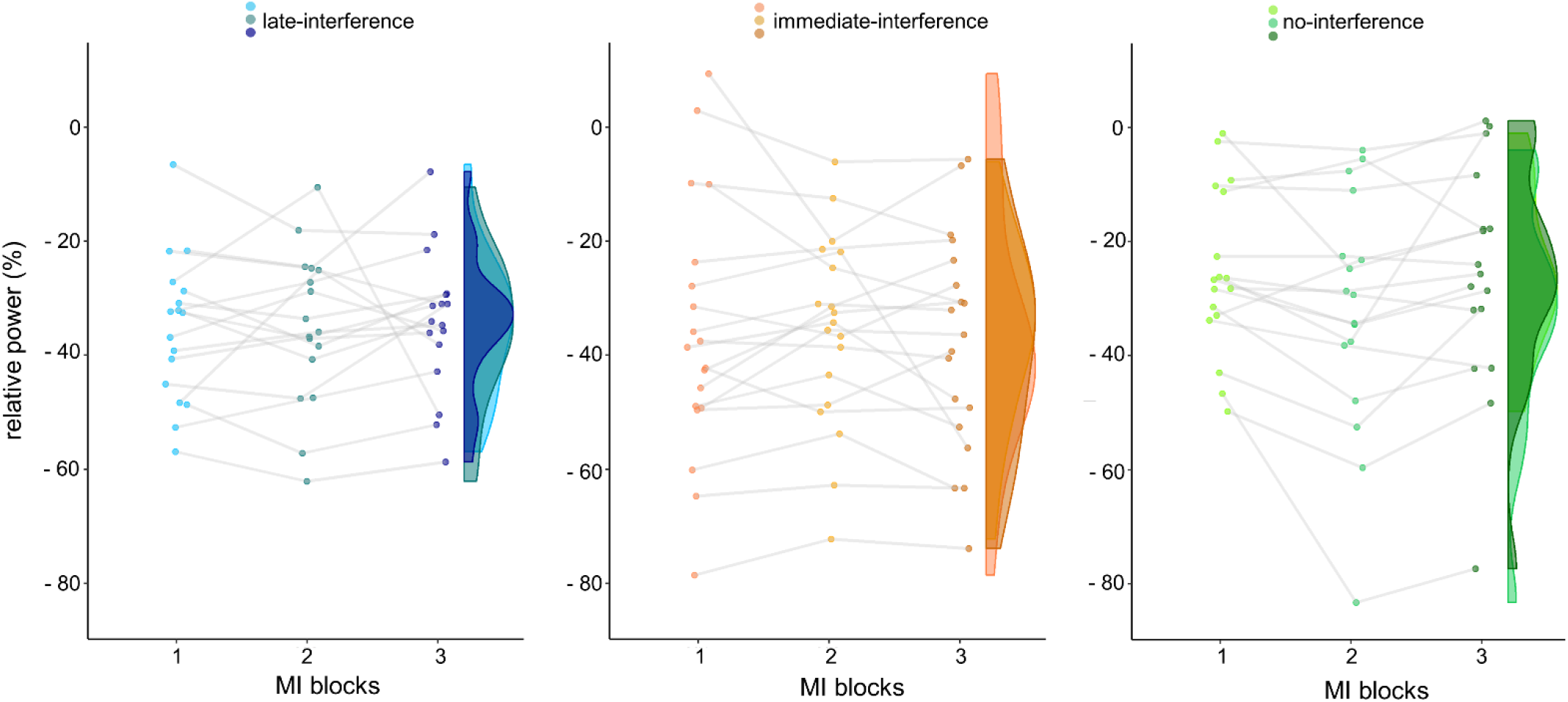
Single subject MI NF ERD data split by group and MI block. Each dot represents the single-subject ERD power value averaged across MI NF runs within a block. Data distribution is estimated using kernel density estimations as implemented in the van Langen open-visualizations repository (van Langen, 2020). Wider sections represent a higher probability that a data point of the population will take on the given value. Lower probability is reflected by narrower sections.

### 3.1 Groups characteristics

Questionnaire scores were compared using Bayesian ANOVAs to ensure comparability across groups. Regarding initial motivation there was moderate evidence^1^ for no difference between groups (BF_group_ = 0.26, error% = 0.04; M ± SD: *late-interference*: 70.12 ± 20.2%; *immediate-interference*: 69.84 ± 18.8%; *no-interference*: 76.65 ± 13.6%). The same held for the initial tiredness level (BF_group_ = 0.26, error% = 0.04; M ± SD: *late-interference*: 41.65 ± 17.7%; *immediate-interference*: 38.26 ± 20.2%; *no-interference*: 33.06 ± 22.4%) also providing moderate evidence for no initial difference between groups. For MI vividness we found anecdotal evidence for no initial difference between groups (BF_group_ = 0.39, error% = 0.02; M ± SD: *late-interference*: 2.97 ± 0.5; *immediate-interference*: 3.13 ± 0.5; *no-interference*: 3.32 ± 0.9) and for MI easiness moderate evidence for no difference between groups (BF_group_ = 0.24, error% = 0.03; M ± SD: *late-interference*: 2.85 ± 0.8; *immediate-interference*: 3.11 ± 0.7; *no-interference*: 3.21 ± 1.1). Further, we found anecdotal evidence for no difference in initial sleepiness scores (SSS; BF_group_ = 0.99, error% = 0.01; M ± SD: *late-interference*: 2.18 ± 0.7; *immediate-interference*: 2.42 ± 0.5; *no-interference*: 2.00 ± 0.4) and moderate evidence for no difference between groups regarding sleep quality (PSQI; BF_group_ = 0.19, error% = 0.03; M ± SD: *late-interference*: 5.47 ± 2.5; *immediate-interference*: 4.90 ± 2.6; *no-interference*: 5.06 ± 1.35).

To ensure comparability between groups regarding initial MI NF performance a 1×3 Bayes ANOVA with between-subject factor group (three levels: *late-interference*, *immediate-interference*, *no-interference*) and MI NF ERD obtained from MI NF block 1 as dependent variable was conducted. The resulting BF of 0.65 suggested anecdotal evidence for H0, that is no initial difference between groups in MI NF ERD.

### 3.2 MI NF gain and MI NF performance over blocks

To examine the effect of NF on MI ERD, we performed paired t-tests within each group and block to compare the training run MI ERD with MI ERD obtained from both MI NF runs within the same block (cf. Fig. 4). For each block and group the MI induced ERD was significantly stronger in the MI NF runs compared to the training run (see Table 1 for details).

**Fig. 4.**
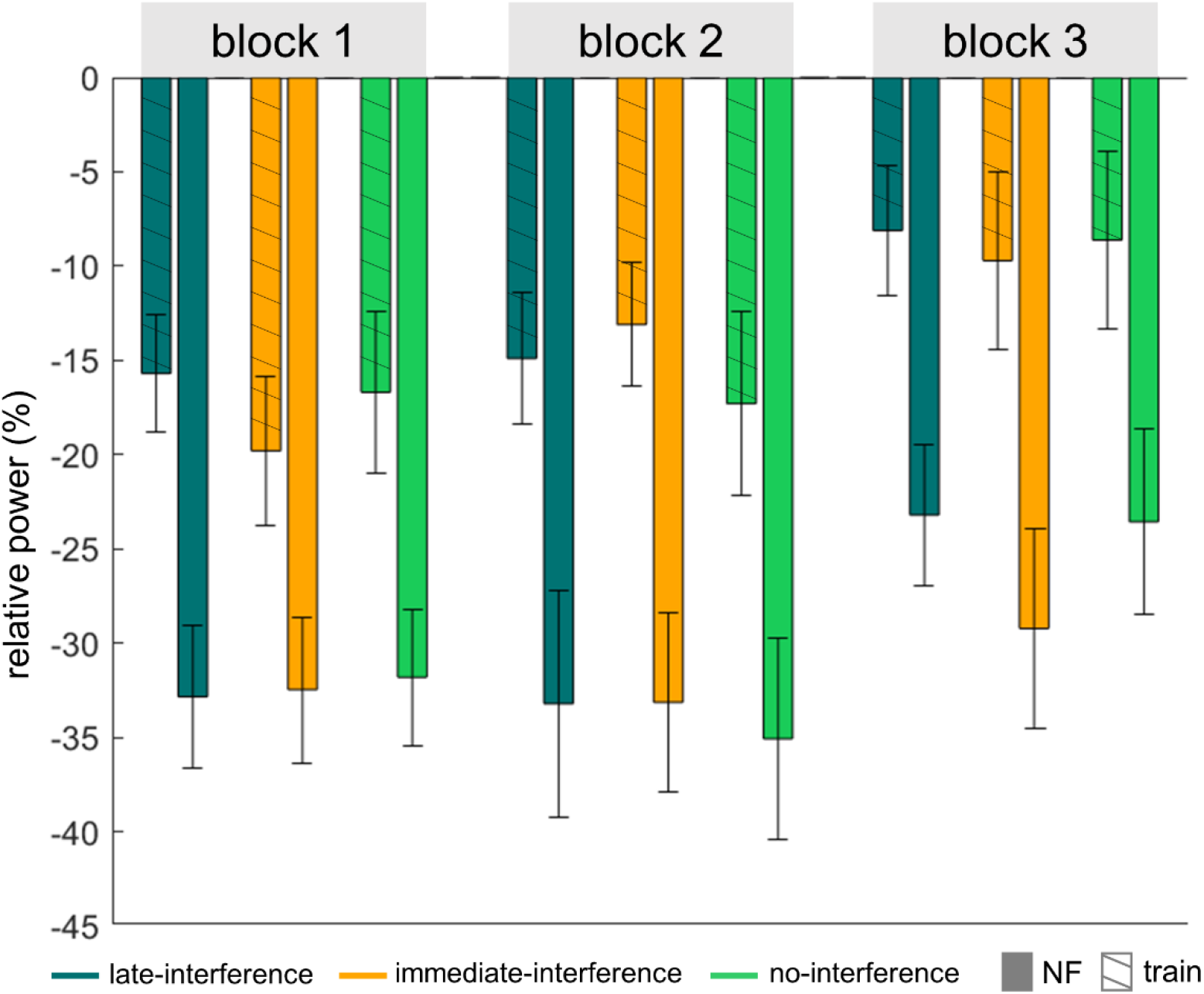
MI ERD for training and NF blocks. Shown are MI (NF) ERDs separate for MI blocks and groups (means ± one standard error). For all combinations of block and group, a strong NF effect is evident in the much stronger ERD in the NF compared to the training runs. However, the NF effect (difference between NF and training runs) is comparable between groups and MI blocks.

**Table 1:** Paired t-tests between training and MI NF runs per block for each group.

| group | blocks | *t* | *p* | *d* | *N* | *M*<sub>train</sub> | *SD*<sub>train</sub> | *M*<sub>NF</sub> | *SD*<sub>NF</sub> |
| --- | --- | --- | --- | --- | --- | --- | --- | --- | --- |
| *late-interference* | 1 | 5.23 | .001 | 1.2 | 17 | -15.7 | 13.63 | -32.85 | 16.34 |
|  | 2 | 3.21 | .008 | 0.74 | 17 | -19.8 | 17.21 | -32.47 | 16.86 |
|  | 3 | 3.1 | .008 | 0.71 | 17 | -16.7 | 18.67 | -31.82 | 15.83 |
| *immediate-interference* | 1 | 3.58 | .002 | 0.78 | 19 | -14.9 | 14.51 | -33.21 | 24.85 |
|  | 2 | 5.67 | <.001 | 1.24 | 19 | -13.1 | 13.45 | -33.14 | 19.59 |
|  | 3 | 4.78 | <.001 | 1.04 | 19 | -17.3 | 20.06 | -35.07 | 21.9 |
| *no-interference* | 1 | 4.9 | <.001 | 1.12 | 17 | -8.12 | 15.13 | -23.2 | 16.43 |
|  | 2 | 4.95 | <.001 | 1.14 | 17 | -9.71 | 20.39 | -29.23 | 23.03 |
|  | 3 | 4.99 | <.001 | 1.15 | 17 | -8.62 | 20.48 | -23.56 | 21.47 |

To explore how this NF gain evolves over blocks and across groups, we performed a 3×3 mixed ANOVA with group as between-subject factor (three levels: *late-interference*, *immediate-interference*, *no-interference*), MI NF block as within-subject factor (three levels: 1, 2, 3) and MI ERD NF gain (i.e. difference between training and NF runs) as dependent variable. We did not find a significant effect (group: F_2,50_ = 0.37, p = .69, η^2^ = 0.02, block: F_1.77,50_ = 0.25, p = .75, η^2^ = 0.01, group x block: F_3.54,50_ = 0.69, p = .58, η^2^ = 0.03). As the ANOVA did not indicate significant effects a Bayes ANOVA was conducted with the same parameters. It provided moderate evidence for no differences between groups and blocks as well as no interaction effects (BF_group_ = 0.22, error% = 0.80; BF_block_ = 0.08, error% = 1.47; BFInclusion = 0.11).

### 3.3 Effect of immediate declarative interference on MI NF ERD

To test whether interference immediately after MI NF practice reduced MI NF ERD over a period of wakefulness, the second MI NF block (MI2) of *groups immediate-interference* and *no-interference* were compared in an independent samples t-test. We did not find a significant effect (*immediate-interference*: M = −35.64%, SD = 16.77%, *no-interference*: M = − 32.01%, SD = 20.77%; t_1,34_ = 0.58, p = 0.57, *d* = 0.19). A Bayes t-test conducted with the same parameters provided further anecdotal evidence for no difference in MI NF block 2 between both groups (BF = 0.37, error% = 0.01). Thus, our data provide no evidence for the hypothesis that immediate interference after MI NF practice reduced MI NF ERD over a period of wakefulness.

There was no evidence for an adverse effect of immediate declarative interference on MI NF ERD. Nonetheless, the planned Bayes paired t-test between MI NF blocks 2 and 3 of *group immediate-interference* was run to confirm that there was also no change after a night of sleep. Results again suggested no difference in MI NF ERD (BF=0.32 (error% = 0.03); moderate evidence for no difference between MI NF blocks 2 and 3).

### 3.4 Early boost and effects of sleep on contralateral MI NF ERD

To test for an early boost effect for MI NF ERD and, if present, its stability over a night of sleep, a 1×3 repeated measures ANOVA with MI NF block (three levels: 1, 2, 3) as within-subject factor and MI NF ERD as dependent variable was performed for *group late-interference*. We did not find a significant effect (F_2,32_ = 0.09, p = 0.91, η^2^ = 0.01). Bayes paired t-tests between MI NF blocks 1 and 2 and between MI NF blocks 2 and 3 provided moderate evidence for no difference between blocks (1 vs 2: BF = 0.25, error% = 0.01; 2 vs. 3: BF = 0.26, error% = 0.01). Hence, our data do not provide evidence for an early boost effect in MI NF ERD nor for a change in MI NF ERD following a night of sleep.

## 4. Discussion

This study investigated the effects of declarative interference and sleep on MI NF performance. Three groups underwent three MI NF blocks each on two consecutive days. Groups differed regarding the presence and timing of a set of declarative interference tasks. We expected an adverse impact of immediate declarative interference on MI NF skill acquisition that would not be recovered by sleep. We further anticipated an early boost for MI NF performance, and if present, we expected sleep to consolidate the MI NF skill at this early boost level. A significant NF effect was present in all three groups and blocks, i.e., ERD was significantly stronger in the NF runs when compared to the training run without NF. Inconsistent with our hypotheses, we found neither evidence for an impact of immediate declarative interference on MI NF ERD nor for an effect of sleep or an early boost effect for MI NF ERD.

Consistent with previous MI NF studies, we found that MI NF resulted in a significantly stronger ERD compared to a training run without NF (see e.g. Darvishi et al., 2017; Zich, Debener, et al., 2015). This feedback effect was present for all groups in all MI blocks, suggesting that task-related feedback reliably enhances MI ERD, independent of the implementation of interference (i.e., immediate, late or no interference) or time of day in this study. However, the MI NF ERD gain did not visibly increase over the course of three MI blocks, which was confirmed by a Bayes analysis suggesting anecdotal evidence for no difference in MI NF ERD gain between blocks and for the absence of an interaction effect of block and interference group. That is, based on the present data, there is neither sufficient evidence for positive practice effects nor for an inhibiting impact of declarative interference for MI NF ERD. This contrasts with the findings of several other studies reporting an increase in MI NF performance with practice (e.g. Foldes, Boninger, Weber, & Collinger, 2020; McWhinney, Tremblay, Boe, & Bardouille, 2018; but see Perronnet et al., 2017; Zich, Debener, et al., 2015). One factor limiting MI NF practise effects in this study may be the break block that consisted of nature documentaries. This break block followed MI blocks in all three groups. Watching nature documentaries is a common control task in (motor) skill acquisition associated studies (Bassolino, Campanella, Bove, Pozzo, & Fadiga, 2014; Friedrich & Beste, 2020; Ruffino, Bourrelier, Papaxanthis, Mourey, & Lebon, 2019; Ruffino et al., 2017) but it is usually not integrated in the experimental group procedure as it is in this study. Thus, it cannot be excluded that the break blocks interfered with the formation of MI NF practice effects. For instance, it is conceivable that participants got distracted by the content of the documentaries or became drowsy and were not able to adequately follow the subsequent MI NF block. A recent study supports the notion that watching a documentary over a prolonged duration as a break occupation or control condition might be problematic. Hachard and colleagues observed that unexpectedly, their control condition of watching a documentary had adverse effects on performance in subsequent balance control tests in young, healthy participants (Hachard, Noé, Ceyte, Trajin, & Paillard, 2020). The authors discuss that participants’ reduced performance in balance control following the control condition could be due to a deleterious effect of prolonged sitting. While this explanation cannot be applied to the present study, an alternative explanation is that watching the documentary induced mental fatigue. This explanation could also hold for the present study However, in the present study, notably, MI NF performance did not deteriorate but remained rather stable between blocks. Thus, no firm conclusions can be drawn but further studies are necessary to systematically study the interplay of MI NF task and control or break tasks.

Immediate declarative interference did not reduce subsequent MI NF ERD when compared to no interference. This result is contrary to our expectation but was statistically confirmed by a Bayes analysis indicating that performance in the second MI block did not differ between a group exposed to declarative interference tasks and a group not exposed to these tasks. To the best of our knowledge, no other study has investigated the impact of declarative interference on MI NF ERD. However, the finding of no effect contrasts with recent studies reporting impaired motor consolidation after MI and ME practice following declarative interference, as reflected in reduced physical task performance (Brown & Robertson, 2007; Debarnot et al., 2012; but see Rothkirch et al., 2018). Reasons for this difference might be closely related or even identical to the reasons leading to another unexpected outcome, i.e. the lack of evidence for an early boost effect on MI NF ERD. Bayes analysis showed moderate evidence for the absence of an early boost effect. Here, as before, a direct comparison to other MI ERD NF studies is not possible due to a lack of published research. But the result clearly contrasts with previous ME and MI practice studies showing early boost effects for both (Debarnot, Clerget, et al., 2011; Nettersheim et al., 2015). One aspect contributing to the deviating results could be the different dependent variables, i.e. MI NF ERD versus physical task performance, and characteristics of these variables, such as test-retest reliability. Another aspect could be unspecific interference through the documentaries used to fill the break block in the present study. If unspecific interference is an issue, it may not only affect MI NF ERD practice effects over time as discussed above, but also the MI NF ERD early boost and specific, declarative interference effects.

An alternative explanation for the discussed dissociation between the findings of previous studies on ME and MI practice and the present one is task difficulty. It is conceivable that our results differ from the aforementioned studies both regarding interference effects and the early boost effect because of the applied motor task. Studies reporting effects of interference and early boost typically concentrate on motor sequence learning tasks with a resulting increase in motor execution data (e.g. Brown & Robertson, 2007; Debarnot et al., 2012; Debarnot, Castellani, et al., 2011). In contrast, we opted for a simple thumb-to-finger-opposition task that has been validated in the context of MI NF for young healthy and older healthy participants (e.g. Nikulin, Hohlefeld, Jacobs, & Curio, 2008; Zich, Debener, Thoene, Chen, & Kranczioch, 2017). This task might however have been too simple to yield measurable behavioural and ERD effects of interference and early boost in healthy young participants. Indeed, the search for MI paradigms suitable for NF research is an ongoing challenge. Research to date is mostly conducted in healthy, often young volunteers. Yet NF interventions, in particular in the neurorehabilitation field, are targeted at clinical, often older populations. Tasks showing practice gains in young and healthy volunteers tend to have little every-day relevance or are conceptualized for a specific audience e.g. athletes (Mulder, Zijlstra, Zijlstra, & Hochstenbach, 2004; Paris-Alemany, La Touche, Gadea-Mateos, Cuenca-Martínez, & Suso-Martí, 2019). Attempts have been made to implement more complex tasks that are both feasible for basic research in non-clinical populations and that have every-day relevance. An example illustrating this is a visuo-motor reach and grasp paradigm with varying levels of difficulty (Allami et al., 2014; Daeglau et al., 2020). Though the visuo-motor reach and grasp paradigm addresses task complexity and generalizability; it brings about new challenges. So do the variable trial durations that result from the setup complicate the calculation and interpretation of the ERD. Furthermore, more complex tasks also mean an increase in cognitive load during MI. This increase reduces cognitive resources available for processing and incorporating NF while doing MI, creating yet another challenge for future research.

Based on experimental groups, MI NF ERD did not change with MI NF practice. In line with this, single subject MI NF ERD shows a remarkable variability over time. Yet nonetheless, visual inspection of MI NF ERDs indicates that all experimental groups contained participants that seemed to not improve at all, participants that improved from Block 1 to Block 2 but lost that gain after a night of sleep, and participants improving continuously over the three blocks. This variation can be seen as indication that individual factors play a major role for MI NF skill acquisition (Ahn & Jun, 2015; Daeglau et al., 2020; Jeunet, Nkaoua, Subramanian, Hachet, & Lotte, 2015; Roc et al., 2019; Zapala, Małkiewicz, Francuz, Kołodziej, & Majkowski, 2019). These individual factors might be more important than the context factors sleep, and declarative interference studied here, or they might interact with them. Future studies should therefore strive to consider individual in addition to context factors instead of looking at both separately.

## 5. Conclusion

The results of the present study suggest that findings of adverse effects of interference or an early boost on motor skill performance cannot readily be generalized to MI NF skill acquisition. As previously shown, the interplay of MI (NF) and performance increase is complex (Dickhaus, Sannelli, Müller, Curio, & Blankertz, 2009; Lotte et al., 2019; Vidaurre & Blankertz, 2010). Context factors are likely contributing to this complexity. In order to better understand their role, it is inevitable to continue to systematically depict common and differing features of MI, MI NF, and ME skill acquisition.

## Data Availability

The datasets generated for this study are available on request to the corresponding author.

## Conflicts of Interest

The authors declare that there is no conflict of interest regarding the publication of this paper.

## Funding Statement

This work was funded by a research grant of the German Research Foundation (Deutsche Forschungsgemeinschaft) [KR3433/3–1]. C.Z. is supported by the Brain Research UK (201718-13, 201617-03).

## Acknowledgments

The authors want to thank Stefan Debener for productive discussions and valuable feedback on an earlier draft of this manuscript, Reiner Emkes for his important contributions to the technical setup, Pauline Roehn and Marius Krösche for their reliable support during the data acquisition phase, Kirsten Netter for her constructive physiotherapeutic advice on EMG recordings, and all participants who volunteered for this study.

## Author contributions

**Mareike Daeglau**: Conceptualization, Formal analysis, Investigation, Methodology, Visualization, Writing—original draft, Writing—review and editing; **Catharina Zich**: Conceptualization, Methodology, Supervision, Writing—review and editing; **Julius Welzel**: Investigation, Writing—review and editing; **Samira Kristina Saak**: Investigation, Writing— review and editing; **Jannik Florian Scheffels**: Investigation, Writing—review and editing; **Cornelia Kranczioch**: Conceptualization, Funding acquisition, Methodology, Project administration, Writing—review and editing.

## Footnotes

1 A Bayes factor (BF) of 1 corresponds to no evidence for either H1 or H0. Bayes factors < 1 indicate that the data provide evidence in favor of H0. Commonly used categories are: BF 1/3-1 ‘anecdotal evidence for H0’; BF 1/10-1/3 ‘moderate evidence for H0’, BF 1/10-1/30 ‘strong evidence for H0’, BF 1/30-1/100 ‘very strong evidence for H0’ and BF < 1/100 ‘extreme evidence for H0’. BF > 1 indicate evidence ranging from ‘anecdotal’ to ‘extreme’ in favor of H1.

## Notes

### Competing Interest Statement

The authors have declared no competing interest.

